# Comparative efficacy of the novel diarylquinoline TBAJ-587 and bedaquiline against a resistant *Rv0678* mutant in a mouse model of tuberculosis

**DOI:** 10.1101/2020.11.17.387977

**Authors:** Jian Xu, Paul J. Converse, Anna M. Upton, Khisimuzi Mdluli, Nader Fotouhi, Eric L. Nuermberger

## Abstract

Since its conditional approval in 2012, bedaquiline (BDQ) has been a valuable tool for treatment of drug-resistant tuberculosis. More recently, a novel short-course regimen combining BDQ with pretomanid and linezolid won approval to treat highly drug-resistant tuberculosis. Clinical reports of emerging BDQ resistance have identified mutations in *Rv0678* that de-repress the expression of the MmpL5/MmpS5 efflux transporter as the most common cause. Because the effect of these mutations on bacterial susceptibility to BDQ is relatively small (e.g., 2-8× MIC shift), increasing the BDQ dose would increase antibacterial activity but also pose potential safety concerns, including QTc prolongation. Substitution of BDQ with another diarylquinoline with superior potency and/or safety has the potential to overcome these limitations. TBAJ-587 has greater *in vitro* potency than BDQ, including against *Rv0678* mutants, and may offer a larger safety margin. Using a mouse model of tuberculosis and different doses of BDQ and TBAJ-587, we found that against wild-type *M. tuberculosis* H37Rv and an isogenic *Rv0678* mutant, TBAJ-587 has greater efficacy against both strains than BDQ, whether alone or in combination with pretomanid and either linezolid or moxifloxacin and pyrazinamide. TBAJ-587 also reduced the emergence of resistance to diarylquinolines and pretomanid.

## INTRODUCTION

Since its conditional approval in late 2012, bedaquiline (BDQ) has become a preferred drug for treatment of multidrug-resistant (MDR) and extensively drug-resistant (XDR) tuberculosis (TB) (1). Mounting clinical evidence confirms the strong bactericidal and sterilizing activity originally observed in mouse models of TB (2–13). More recently, novel regimens based on the backbone of BDQ and pretomanid (PMD) have shown the potential to significantly shorten the duration of treatment of MDR/XDR-TB as well as drug-susceptible (DS) TB (14, 15), as predicted by studies in mice (16, 17). Specifically, a BDQ, PMD and linezolid (LZD) regimen (also abbreviated as BPaL) resulted in an unprecedented 90% rate of successful outcomes in XDR-TB and difficult-to-treat MDR-TB patients with just 6 months of treatment in the Nix-TB trial (14); and in the NC-005 trial, the combination of BDQ, PMD, moxifloxacin (MXF) and pyrazinamide (PZA) (also abbreviated as BPaMZ) achieved faster sputum culture conversion among MDR-TB patients than the first-line standard-of-care regimen (SOC) did in DS-TB patients (15). The ongoing SimpliciTB trial (ClinicalTrials.gov Identifier: NCT03338621) is evaluating a 4-month duration of BPaMZ against the 6-month SOC regimen in DS-TB.

Despite the positive clinical outcomes observed thus far with BDQ-containing regimens, opportunities for further optimization exist. In one pharmacometric analysis, the median average continuation phase plasma BDQ concentration was below the level associated with 50% of its maximal effect (EC_50_) in MDR-TB patients receiving the registered dose, indicating that higher BDQ doses would achieve greater efficacy (15). However, potential safety concerns, including QTc prolongation, have limited enthusiasm for testing higher BDQ doses (18). Evidence also continues to emerge that the efficacy of BDQ-containing regimens can be compromised by inactivating mutations in the *Rv0678* gene, which encodes a negative transcriptional regulator of the *mmpL5-mmpS5* transporter in *Mycobacterium tuberculosis* (19–22). Although mutations in *Rv0678* are associated with relatively small reductions in susceptibility to BDQ and clofazimine, they are readily selected by BDQ and/or clofazimine treatment in mouse models of TB and also have been selected during clinical use of BDQ, including in the Nix-TB trial (14). Coupled with evidence that *Rv0678* variants with reduced BDQ susceptibility have been isolated from MDR-TB patients without known prior exposure to BDQ or clofazimine (19, 23, 24), these reports raise concern that emergence of *Rv0678* variants could undermine the promising clinical efficacy of BDQ-containing regimens.

Development of diarylquinoline drugs with superior potency could mitigate the aforementioned concerns associated with BDQ. For example, TBAJ-587 has greater *in vitro* potency than BDQ and reduced cardiovascular liability (25). Although mutation of *Rv0678* reduces the *in vitro* activity of TBAJ-587 to a similar degree as BDQ, TBAJ-587 remains more potent than BDQ against such mutants and therefore may be more effective at killing them or preventing their selection during treatment (26).

In the current experiment, we evaluated the dose-dependent effects of BDQ (B) and TBAJ-587 (S) against an *Rv0678* loss-of-function mutant compared to the wild-type H37Rv parent, to determine the impact of such mutations on the activity of BDQ and its contribution to the efficacy of the BPaL and BPaMZ regimens. The results indicate that replacing BDQ in these regimens with TBAJ-587 could increase their efficacy against wild-type *M. tuberculosis* as well as *Rv0678* mutants and reduce the emergence of resistance to diarylquinolines and companion drugs.

## RESULTS

### Bacterial strains and mouse infection model

The experimental schemes indicating the regimens used against the wild-type strain, *M. tuberculosis* H37Rv, and an isogenic mutant with an IS6110 insertion in the *Rv0678* gene are in Supplementary Tables 1 and 2, respectively. Using the broth macrodilution method in 7H9 media, the MICs of BDQ and TBAJ-587 against the H37Rv strain were 0.0625 and 0.016, respectively, while against the *Rv0678* mutant, the MICs were 0.5 and 0.0625, respectively.

One day after high-dose aerosol infection of BALB/c mice with either strain, mean (±S.D.) lung CFU counts were 4.11±0.06 log_10_ for H37Rv and 4.19±0.08 for the *Rv0678* mutant. Mean CFU counts increased over the following two weeks to 7.79±0.11 and 7.75±0.10, respectively, when treatment began (Table 1).

**Table 1.**
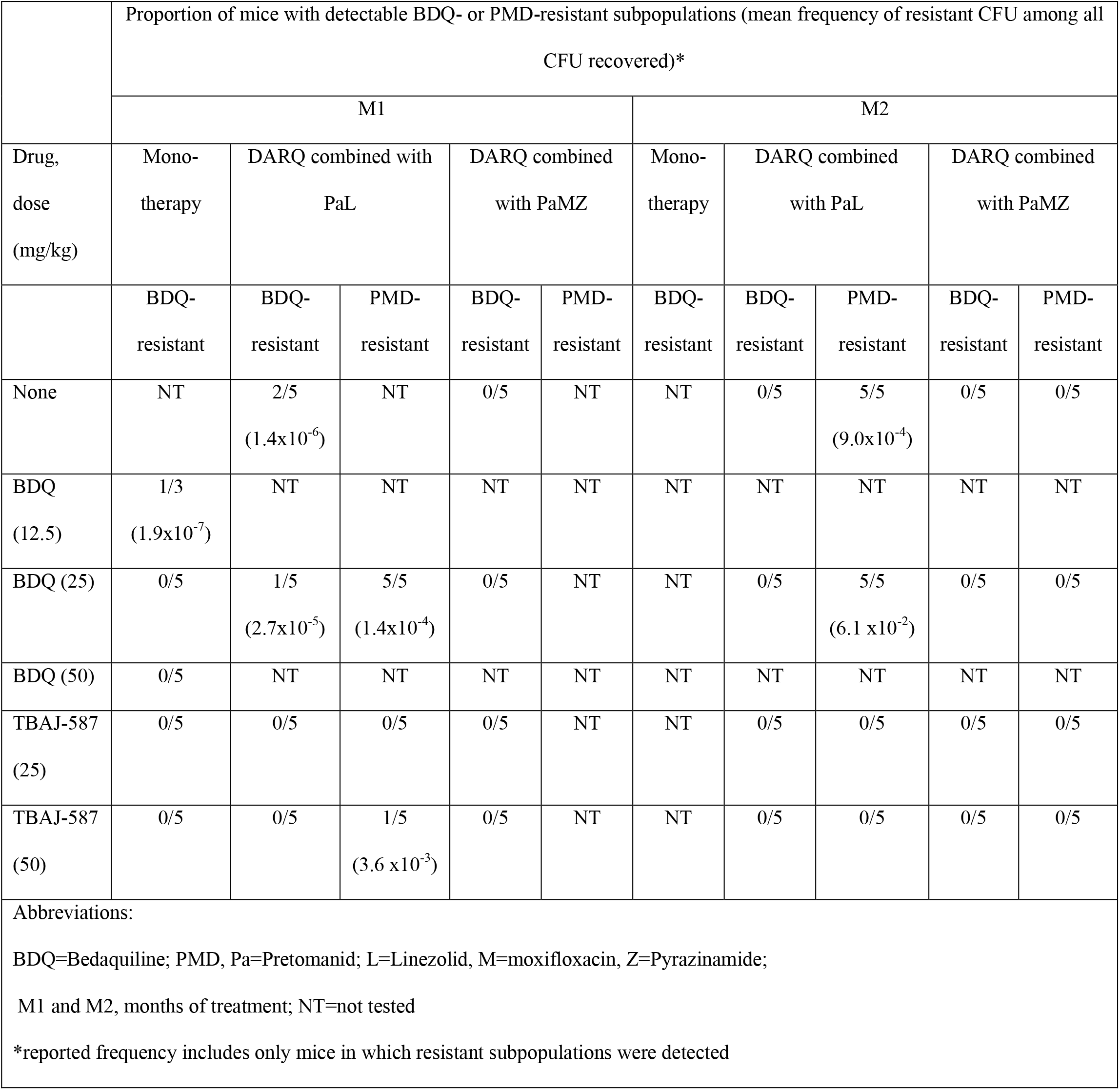
Proportion of mice showing resistance to bedaquiline (BDQ) and pretomanid (PMD) after infection with wild-type *M. tuberculosis* and antimicrobial treatment.

### Response to Treatment. (i) Mice infected with wild-type *M. tuberculosis* H37Rv

As shown in Fig. 1A and Supplementary Table 3, after one month of treatment, BDQ at 25 mg/kg alone reduced the mean lung burden by 3.29 log_10_ compared to Day 0 (D0) and was more active than the combination of PMD and LZD. When added to PMD and LZD, BDQ reduced the mean CFU count by an additional 2.88 log_10_ (i.e., BPaL vs. PaL). The combination with PMD, MXF, and PZA was much more active than PMD plus LZD, and the addition of BDQ reduced the mean CFU count by an additional 1.77 log_10_ (i.e., BPaMZ vs. PaMZ). Treatment with TBAJ-587 at 25 and 50 mg/kg resulted in reductions of lung CFU burden that were more than 1.5 log_10_ greater than observed with BDQ alone (p<0.0001). There was no statistically significant difference between TBAJ-587 at 25 mg/kg compared to 50 mg/kg. When combined with PMD and LZD, the effect of TBAJ-587 again reduced the mean CFU burden by >1.5 log_10_ compared to BDQ. The S_25_PaL regimen was significantly more active than B_25_PaL (p=0.0002) and not significantly less active than S_50_PaL (p=0.0954). Similarly, in combination with PMD, MXF, and PZA, TBAJ-587 reduced the mean CFU count by approximately 1.5 log_10_ more than BDQ did. The S_25_PaMZ regimen was significantly more active than B_25_PaMZ (p=0.0001) and not significantly different from S_50_PaMZ (p=0.9049). The response to monotherapy was also evaluated at Month 2 (Fig. 1B) and showed that both diarylquinolines continued to reduce the lung CFU burden but that TBAJ-587 continued to kill at a faster rate than BDQ.

**FIG 1.**
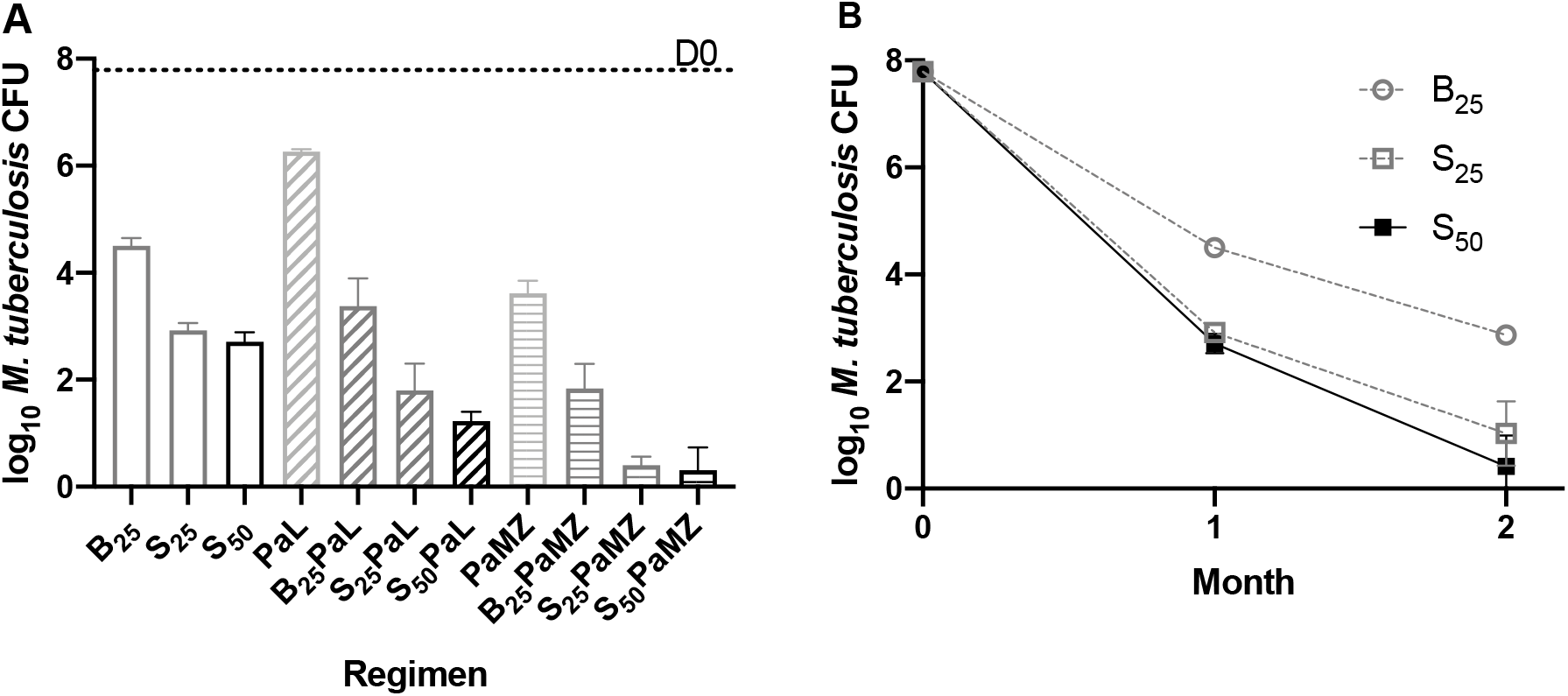
TBAJ-587 (S) is more active than bedaquiline (B) either alone or in combination with pretomanid and linezolid (PaL) or pretomanid, moxifloxacin and pyrazinamide (PaMZ) against wild-type *M. tuberculosis*. (A) Activity of different regimens during the first month of treatment. (B) Activity of diarylquinoline monotherapy during two months of treatment.

### Response to Treatment. (ii) Mice infected with *M. tuberculosis* H37Rv with an *Rv0678* mutation

As shown in Fig. 2A and Supplementary Table 4, dose-dependent reductions in lung CFU were observed with both BDQ (at 12.5, 25, and 50 mg/kg) and TBAJ-587 (at 25 and 50 mg/kg) monotherapy. Both DARQs were significantly less active against the mutant than against the wild-type H37Rv strain, but a similar difference in potency was observed between BDQ and TBAJ-587 against each strain. BDQ at 25 mg/kg reduced the mean CFU count by only 0.56 log_10_ compared to D0. TBAJ-587 at 25 mg/kg reduced the mean CFU count by 2 log_10_ and was significantly more active than BDQ at any dose (p<0.0001) but less active (p=0.0269) than TBAJ-587 at 50 mg/kg. PMD in combination with either LZD or MXF and PZA had similar activity against each strain. Both BDQ and TBA-587 significantly (p<0.0001) increased the activity of the PaL and PaMZ combinations at M1. As expected, both BDQ and TBAJ-587 contributed a smaller effect size in combination with PaL against the *Rv0678* mutant compared to that against the H37Rv strain. Interestingly, BDQ, and for the most part TBAJ-587, contributed similar effects to the PaMZ combination irrespective of the infecting strain. SPaL and SPaMZ were more active than the corresponding B-containing regimens (p<0.0001 for both doses of S in the SPaL regimens compared to BPaL at M1 and M2 and p=0.0063 and 0.0002 for the SPaMZ regimens compared to BPaMZ at M1 but no difference at M2). A statistically significant difference (p=0.0351) between the TBAJ-587 dose levels in combination with PaMZ but not PaL at M1 was noted. At M2, both diarylquinolines again added significant activity to PaL (p<0.0001). At this time point, TBAJ-587, at either 25 or 50 mg/kg significantly (p=0.0364) increased the activity of PaMZ whereas BDQ did not (p=0.1916).

**FIG 2.**
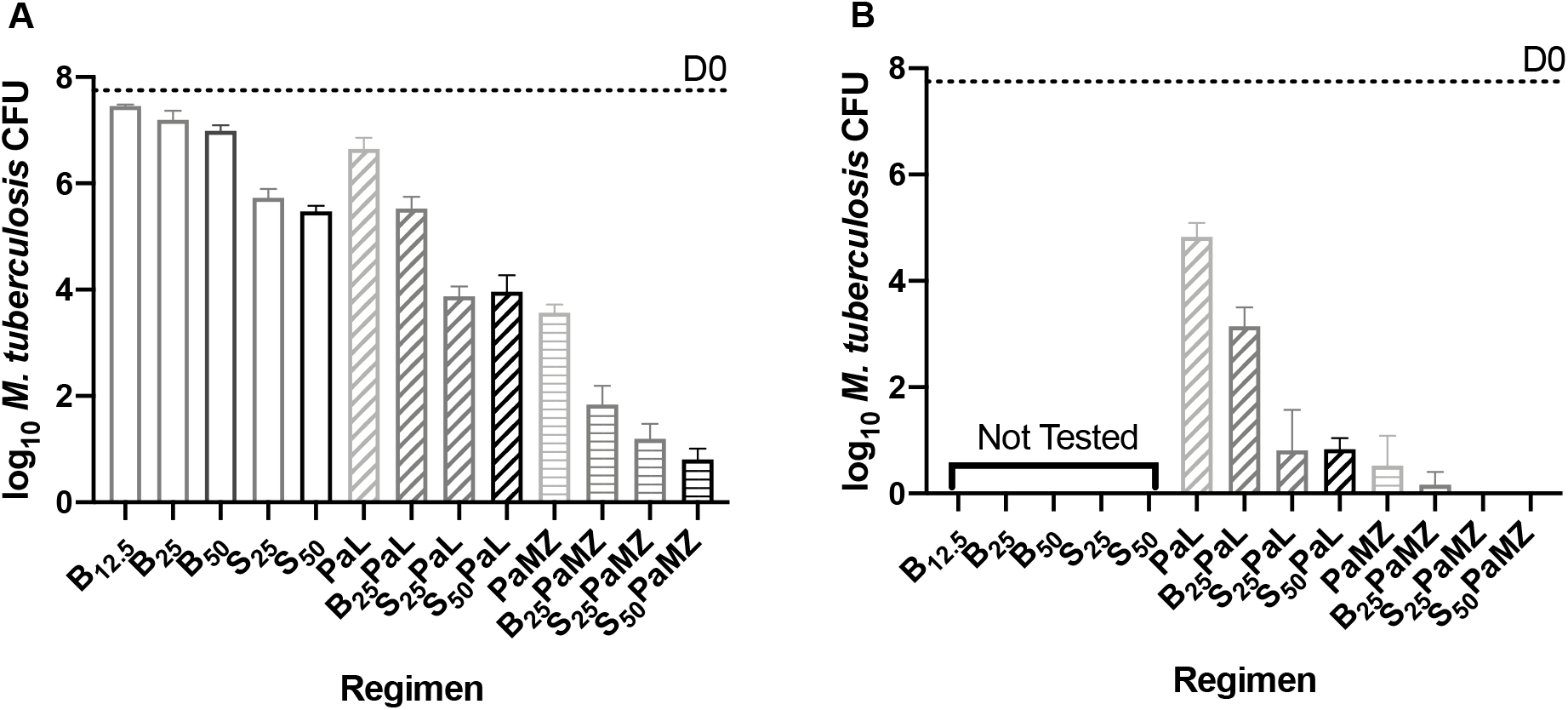
TBAJ-587 (S) is more active than bedaquiline (B) either alone or in combination with pretomanid and linezolid (PaL) or pretomanid, moxifloxacin and pyrazinamide (PaMZ) against *M. tuberculosis* with an *Rv0678* mutation. Activity of the indicated regimens at M1 (A) and M2 (B).

### Selection of drug-resistant mutants. (i) Mice infected with wild-type *M. tuberculosis* H37Rv

At D0, the mean frequencies of CFU able to grow on agar containing 0.06 and 0.25 μg/ml of BDQ were 4.2×10^−5^ and 1.8×10^−5^, respectively. Plates containing BDQ 0.06 μg/ml were used to quantify the resistant subpopulation at subsequent time points. All mice infected with *M. tuberculosis* H37Rv and treated with BDQ monotherapy at 25 mg/kg for one or two months demonstrated selective amplification of BDQ-resistant CFU, which represented approximately 3% and 18% of the recovered CFU after one and two months, respectively (Table 1). However, it should be noted that the total number of CFU isolated on BDQ-containing plates decreased from D0 to M1 to M2, indicating that BDQ treatment reduced the size of the subpopulation of CFU able to grow on BDQ but not nearly as fast as it reduced the number of fully susceptible CFU. Monotherapy with TBAJ-587 at either 25 or 50 mg/kg prevented the selection of spontaneous BDQ-resistant mutants, with the exception of a single colony recovered from one of five mice after two months of TBAJ-587 monotherapy at 25 mg/kg (representing an estimated one-third of all CFU recovered). As expected, the BPaL and BPaMZ regimens resulted in less selection of BDQ-resistant CFU compared to BDQ monotherapy. Only three mice and one mouse receiving BPaL and BPaMZ, respectively, for one month harbored BDQ-resistant CFU. Replacing BDQ with TBAJ-587 at either 25 or 50 mg/kg in combination with PaL or PaMZ prevented the selection of spontaneous BDQ-resistant mutants. As expected, each diarylquinoline prevented selection of PMD-resistant mutants when combined with PaL.

### Selection of drug-resistant mutants. (ii) Mice infected with *M. tuberculosis* H37Rv with an *Rv0678* mutation

Among mice infected with the *Rv0678* mutant (Table 2), the selection of CFU with a higher level of BDQ resistance (as quantified by growth on agar containing BDQ at 1 μg/ml) was rare, being observed as a very low percentage of recovered CFU in only 2 of 5 mice (average frequency of 1.4×10^−6^ CFU) treated with PaL for one month, 1 of 3 surviving mice (frequency of 1.9×10^−7^ CFU) treated with BDQ 12.5 mg/kg for one month, and 1 of 5 mice (frequency of 2.7×10^−5^ CFU) treated with B_25_PaL for one month. No CFU resistant to BDQ at 1 μg/ml were recovered from mice treated with TBAJ-587 alone or in combination for 1-2 months or from mice treated with BDQ alone or in combination for 2 months. Selection of PMD-resistant mutants was more evident among mice infected with the *Rv0678* mutant compared to those infected with the wild-type H37Rv and was associated with receipt of BDQ rather than TBAJ-587. PMD-resistant mutants were recovered from 10 of 10 mice receiving BPaL for 1-2 months and comprised over 6% of bacterial population after two months of treatment. In contrast, among 20 mice receiving SPaL for 1-2 months, only 1 mouse harbored PMD-resistant CFU at a frequency of 0.36%, indicating that TBAJ-587 more effectively eradicated double mutants with resistance to both BDQ (via the baseline *Rv0678* mutation) and PMD (via spontaneous resistance mutations).

**Table 2.**
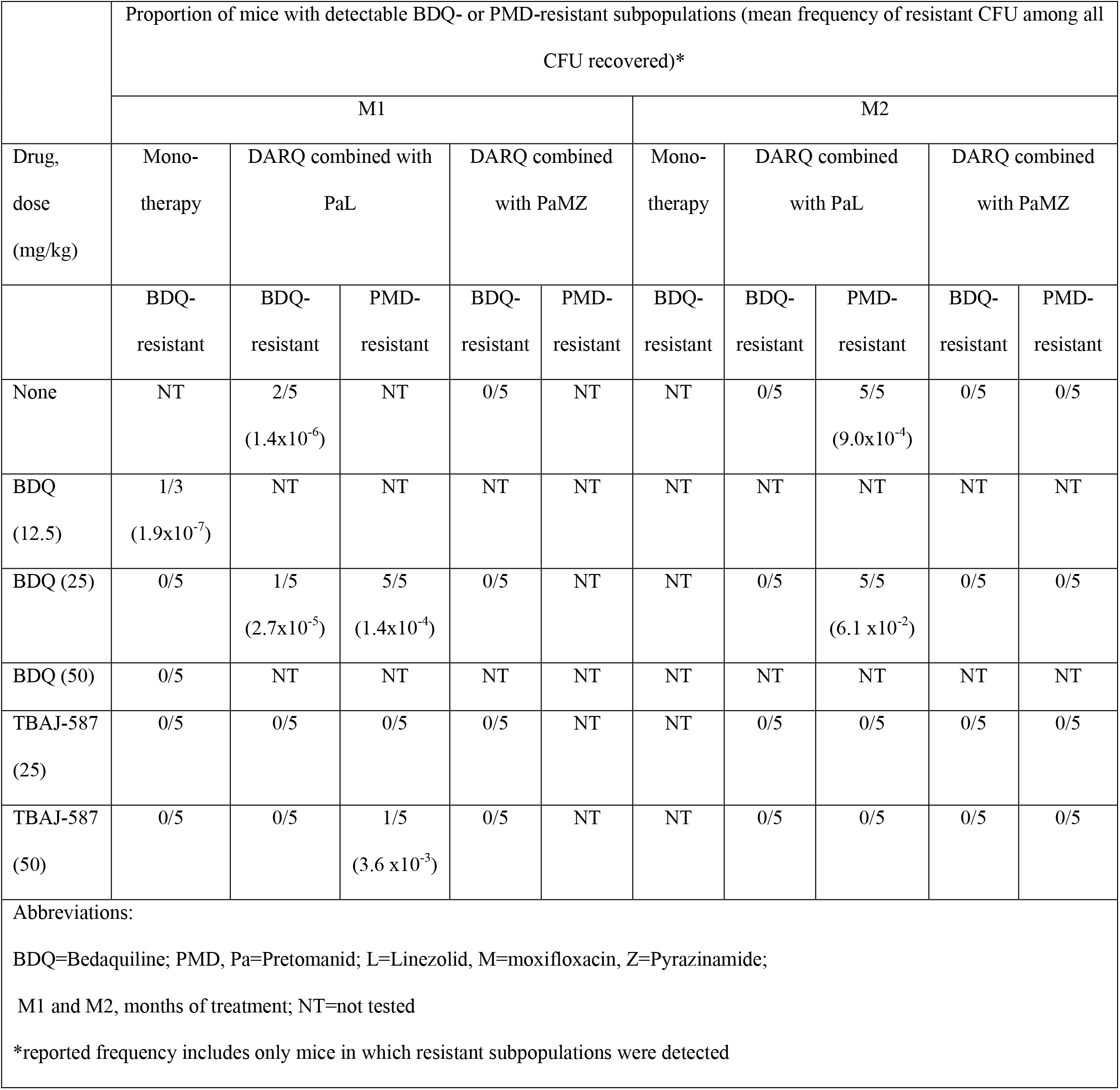
Proportion of mice showing resistance to bedaquiline (BDQ) and pretomanid (PMD) after infection with *M. tuberculosis* containing a *Rv0678* mutation and antimicrobial treatment

## DISCUSSION

The BPaL regimen is now an important fully oral, short-course treatment option for XDR-TB and difficult-to-treat MDR-TB (1, 14). BPaMZ has demonstrated potential to be an even shorter-course regimen for selected MDR-TB and possibly DS-TB (15). Therefore, it is of great concern that *Rv0678* variants with reduced susceptibility to BDQ are reported with increasing frequency among patients treated with BDQ or clofazimine, including at least 1 participant treated with BPaL in the Nix-TB trial who relapsed with an *Rv0678* mutant (14). In addition, several studies have reported *Rv0678* variants among baseline MDR-TB isolates obtained prior to any known BDQ or clofazimine exposure (19, 23, 24), suggesting that they may be enriched among MDR-TB isolates due to selection by other factors. The results presented here confirm that loss-of-function mutations in *Rv0678* significantly reduce the efficacy of BDQ *in vivo* (19) and show for the first time that they also significantly reduced the efficacy of the novel BPaL regimen. Mice infected with the *Rv0678* mutant required two months of BPaL treatment to reach the same CFU count as observed in wild-type infected mice treated for one month. Moreover, spontaneous BDQ-resistant mutants were selectively amplified by BPaL treatment in wild-type-infected mice, approaching or surpassing the 1% threshold commonly used to define resistance by agar proportion method in 3 of 5 mice. Interestingly, BDQ resistance was also observed in one of five mice treated with BPaMZ for one month. Therefore, although the likelihood of BDQ resistance emerging during treatment is undoubtedly higher when BDQ is combined with less effective companion drugs, it should be recognized that BDQ produces strong selective pressure favoring amplification of spontaneous BDQ-resistant mutants even in these highly active regimens.

Inadvertent treatment of patients infected with an *Rv0678* mutant, perhaps even as a heteroresistant subpopulation, could lead to selection of additional mutations conferring resistance to companion drugs. In the present study, dual BDQ and PMD resistance emerged in all 5 mice infected with the *Rv0678* mutant and treated with BPaL for two months. In stark contrast, no PMD-resistant mutants were isolated from wild-type-infected mice treated with BPaL. These results raise concerns that inadvertent treatment of patients infected with an *Rv0678* mutant with BPaL could lead to dangerous new form of multidrug resistance defined by resistance to BDQ and PMD, which would likely extend to delamanid (27, 28).

Surprisingly, the BPaMZ regimen, like the PaMZ regimen, had similar bactericidal effects against both the wild-type and mutant infections, indicating that the contribution of BDQ to the efficacy of BPaMZ was not affected by the *Rv0678* mutation. While further study is needed to confirm this observation and explore its potential mechanism, it is conceivable that PZA reduces the function of the MmpL5/S5 transporter through disruption of membrane potential (29) or has other synergies with BDQ that enable bactericidal effects at lower intrabacillary BDQ concentrations.

In the present study, replacement of BDQ with the more potent diarylquinoline TBAJ-587 improved the bactericidal activity of the BPaL and BPaMZ regimens against the wild-type H37Rv strain, indicating its potential to shorten the duration of treatment needed to prevent relapse (30). The present work also demonstrates that TBAJ-587 is more effective than BDQ against an isogenic *Rv0678* mutant. Although loss of *Rv0678* function causes a similar shift in susceptibility to BDQ and TBAJ-587, TBAJ-587 retains superior potency. At 25 mg/kg/day, BDQ loses most of its bactericidal activity against the *Rv0678* mutant, whereas TBAJ-587 exhibits bactericidal activity (i.e., 2 log kill over 1 month) as monotherapy. Indeed, SPaL was practically as effective against the mutant as BPaL was against the wild-type H37Rv strain. Interestingly, replacing BDQ with TBAJ-587 had the smallest apparent benefit in the BPaMZ regimen against the *Rv0678* mutant.

As a function of its superior activity against *Rv0678* mutants, TBAJ-587 more effectively prevented the emergence of new drug resistance more effectively than BDQ. In mice infected with the wild-type H37Rv strain, TBAJ-587, both alone and in combination with PaL or PaMZ, nearly abolished the selective amplification of spontaneous BDQ-resistant CFU. Whereas BDQ resistance emerged in 100%, 60% and 20% of mice treated with BDQ alone, BPaL and BPaMZ, respectively, it was observed in only 5% of mice treated with TBAJ-587 alone and in none of the the 10 mice each receiving SPaL or SPaMZ. Taken together, these results indicate that, in addition to improving efficacy, replacing BDQ with TBAJ-587 would make regimens like BPaL and BPaMZ more robust to the emergence of diarylquinoline resistance. Importantly, we did not observe selection of higher level, or “second-step” BDQ resistance despite treating the *Rv0678* mutant infection with BDQ or TBAJ-587 alone. This is likely a function of the low frequency of viable spontaneous *atpE* mutants and their fitness costs observed *in vivo* (31). Although *atpE* mutations have been identified in a small number of BDQ-resistant clinical isolates to date (32, 33), *Rv0678* mutations have been more prevalent. Therefore, overcoming *Rv0678*-mediated resistance with more potent diarylquinolines like TBAJ-587 could greatly extend the utility and longevity of this important new class of drugs.

Use of TBAJ-587 in place of BDQ in the BPaL regimen in mice infected with the *Rv0678* mutant also largely prevented the selection of spontaneous PMD-resistant double mutants. Following two months of treatment, all five BPaL-treated mice harbored dual BDQ/PMD-resistant isolates, compared with zero of ten SPaL-treated mice. In fact, BPaL and PaL resulted in similar absolute numbers of dual BDQ/PMD-resistant mutants at the end of 2 months of treatment, indicating that BDQ did not contribute to faster elimination of spontaneous PMD-resistant mutants in the *Rv0678* mutant background. Thus, the benefits of replacing BDQ with TBAJ-587 in the BPaL regimen are expected to include the prevention of PMD resistance (i.e., a new form of multidrug resistance to diarylquinolines and nitroimidazoles) when infection with an *Rv0678* mutant is inadvertently treated with a regimen combining a diarylquinoline with PaL.

## MATERIALS AND METHODS

### Bacterial Strains

The laboratory strain, *M. tuberculosis* H37Rv, and a spontaneous bedaquiline-resistant mutant with an IS*6110* insertion in *Rv0678* at aa116/nt349 were used in this study. The *Rv0678* mutant was previously identified as BDQ-8 when it was isolated from an untreated mouse infected by H37Rv (31). The *Rv0678* mutation and the absence of other mutations in genes associated with drug resistance were confirmed by whole genome sequencing. The MICs of BDQ and TBAJ-587 against both strains were determined by the broth macrodilution method in complete 7H9 broth using polystyrene tubes.

### Infection model

Female BALB/c mice, 6 wks old, were aerosol-infected with ~4 log_10_ CFU of each *M. tuberculosis* strain from a log phase culture with OD_600_ of ~0.8 on D-14. Treatment started 2 weeks later (D0). Mice were sacrificed for lung CFU counts on D-13 and D0 to determine the number of CFU implanted and the number present at the start of treatment, respectively.

### Antibiotic treatment

Mice were treated as indicated in supplementary tables 1 and 2 at the following doses (mg/kg): BDQ (12.5, 25, or 50), TBAJ-587 (25 or 50), PMD (100), MXF (100), LZD (100), and PZA (150) once daily five days per week for one or two months with the exception of BDQ at 50 mg/kg that was given twice daily at 25 mg/kg. BDQ and TBAJ-587 were formulated in 20% HPCD solution acidified with 1.5% 1N HCl. PMD was prepared in the CM-2 formulation as previously described (34). LZD was prepared in 0.5% methylcellulose. MXF and PZA were prepared in water. Except as noted for BDQ at 50 mg/kg, the diarylquinolines and PMD were administered in a single gavage in the morning and LZD, MXF, and PZA were administered in the afternoon.

### Evaluation of drug efficacy

Efficacy determinations were based on lung CFU counts after 1 month and 2 months of treatment. At each time point, lungs were removed aseptically and homogenized in 2.5 ml PBS. Lung homogenates were plated in serial dilutions on 0.4% charcoal-supplemented 7H11 agar with 2 selective antibiotics, i.e., to compensate for charcoal absorption of these drugs. Final concentrations in μg/ml for these antibiotics were: cycloheximide (20), carbenicillin (100), polymyxin B (50), and trimethoprim (40).

### Evaluation of resistance selection

At each time point, lung homogenates from mice infected with the wild-type H37Rv strain were plated in parallel on drug-free 7H11 plates and on the same plates supplemented with 0.06 μg/ml of BDQ. Aliquots of lung homogenates from mice infected with the *Rv0678* mutant were plated on plates containing a higher BDQ concentration (1 μg/ml) to evaluate for *atpE*-mediated resistance (31) and on plates containing 2 μg/ml of PMD.

### Statistical analysis

Differences between regimens were assessed by one-way ANOVA with Dunnett’s multiple comparison correction using GraphPad Prism version 8.

## ACKNOWLEDGEMENTS

This research was supported by TB Alliance with support from Australia Aid, the Bill and Melinda Gates Foundation, the Germany Federal Ministry of Education and Research through KfW, Global Health Innovative Technology Fund, Irish Aid, Netherlands Ministry of Foreign Affairs, UK Aid and the UK Department of Health.

---

**Supplementary Table 1.** Experimental scheme for mice infected with wild-type *M. tuberculosis* H37Rv

| Regimen | No. of mice<sup>a</sup> |  |  |  | Total |
| --- | --- | --- | --- | --- | --- |
|  | D-13 | D0 | M1 | M2 |  |
| Untreated | 5 | 5 |  |  | 10 |
| B<sub>25</sub> |  |  | 5 | 5 | 10 |
| S<sub>25</sub> |  |  | 5 | 5 | 10 |
| S<sub>50</sub> |  |  | 5 | 5 | 10 |
| PaL |  |  | 5 |  | 5 |
| PaMZ |  |  | 5 |  | 5 |
| B<sub>25</sub>PaL |  |  | 5 |  | 5 |
| B<sub>25</sub>PaMZ |  |  | 5 |  | 5 |
| S<sub>25</sub>PaL |  |  | 5 |  | 5 |
| S<sub>25</sub>PaMZ |  |  | 5 |  | 5 |
| S<sub>50</sub>PaL |  |  | 5 |  | 5 |
| S<sub>50</sub>PaMZ |  |  | 5 |  | 5 |
| Total | 5 | 5 | 55 | 15 | 80 |
| <sup>a</sup>Time points shown as days (D-13 or D) or months (M1 or M2) of treatment.<br>Abbreviations:<br>B=Bedaquiline; Pa=Pretomanid; L=Linezolid, M=moxifloxacin,<br>Z=Pyrazinamide; S= TBAJ-587. Doses are indicated in subscripts. |  |  |  |  |  |

**Supplementary Table 2.** Experimental scheme for mice infected with *M. tuberculosis* H37Rv *Rv0678* mutant

| Regimen | No. of mice sacrificed<sup>a</sup> |  |  |  | Total |
| --- | --- | --- | --- | --- | --- |
|  | D-13 | D0 | M1 | M2 |  |
| untreated | 5 | 5 | 5 |  | 15 |
| B<sub>12.5</sub> |  |  | 5 |  | 5 |
| B<sub>25</sub> |  |  | 5 |  | 5 |
| B<sub>50</sub> |  |  | 5 |  | 5 |
| S<sub>25</sub> |  |  | 5 |  | 5 |
| S<sub>50</sub> |  |  | 5 |  | 5 |
| Pa<sub>100</sub>L<sub>100</sub> |  |  | 5 | 5 | 10 |
| PaM<sub>100</sub>Z<sub>150</sub> |  |  | 5 | 5 | 10 |
| B<sub>25</sub>PaL |  |  | 5 | 5 | 10 |
| B<sub>25</sub>PaMZ |  |  | 5 | 5 | 10 |
| S<sub>25</sub>PaL |  |  | 5 | 5 | 10 |
| S<sub>25</sub>PaMZ |  |  | 5 | 5 | 10 |
| S<sub>50</sub>PaL |  |  | 5 | 5 | 10 |
| S<sub>50</sub>PaMZ |  |  | 5 | 5 | 10 |
| Total | 5 | 5 | 70 | 40 | 120 |
| <sup>a</sup>Time points shown as days (D-13 or D) or months (M1 or M2) of treatment.<br>Abbreviations:<br>B=Bedaquiline; Pa=Pretomanid; L=Linezolid, M=moxifloxacin,<br>Z=Pyrazinamide; S= TBAJ-587. Doses are indicated in subscripts. |  |  |  |  |  |

**Supplementary Table 3.** Lung CFU counts assessed during treatment against the *M. tuberculosis* H37Rv strain.

| Regimen | Mean (±SD) log<sub>10</sub> CFU count at<sup>a</sup>: | | | |
| --- | --- | --- | --- | --- |
|  | D-13 | D0 | M1 | M2 |
| Untreated | 4.11±0.06 | 7.79±0.11 | | |
| B<sub>25</sub> | | | 4.50±0.14 | 2.87±0.03 |
| S<sub>25</sub> | | | 2.92±0.14 | 1.03±0.60 |
| S<sub>50</sub> | | | 2.71±0.18 | 0.42±0.57 |
| PaL | | | 6.26±0.05 | |
| B<sub>25</sub>PaL | | | 3.38±0.52 | |
| S<sub>25</sub>PaL | | | 1.80±0.50 | |
| S<sub>50</sub>PaL | | | 1.23±0.18 | |
| PaMZ | | | 3.61±0.23 | |
| B<sub>25</sub>PaMZ | | | 1.84±0.46 | |
| S<sub>25</sub>PaMZ | | | 0.40±0.16 | |
| S<sub>50</sub>PaMZ | | | 0.31±0.43 | |
| <sup>a</sup>Time points shown as days (D-13 or D) or months (M1 or M2) of treatment.<br>Abbreviations:<br>B=Bedaquiline; Pa=Pretomanid; L=Linezolid, M=moxifloxacin, Z=Pyrazinamide; S=TBAJ-587. Doses are indicated in subscripts. |  |  |  |  |

**Supplementary Table 4.**
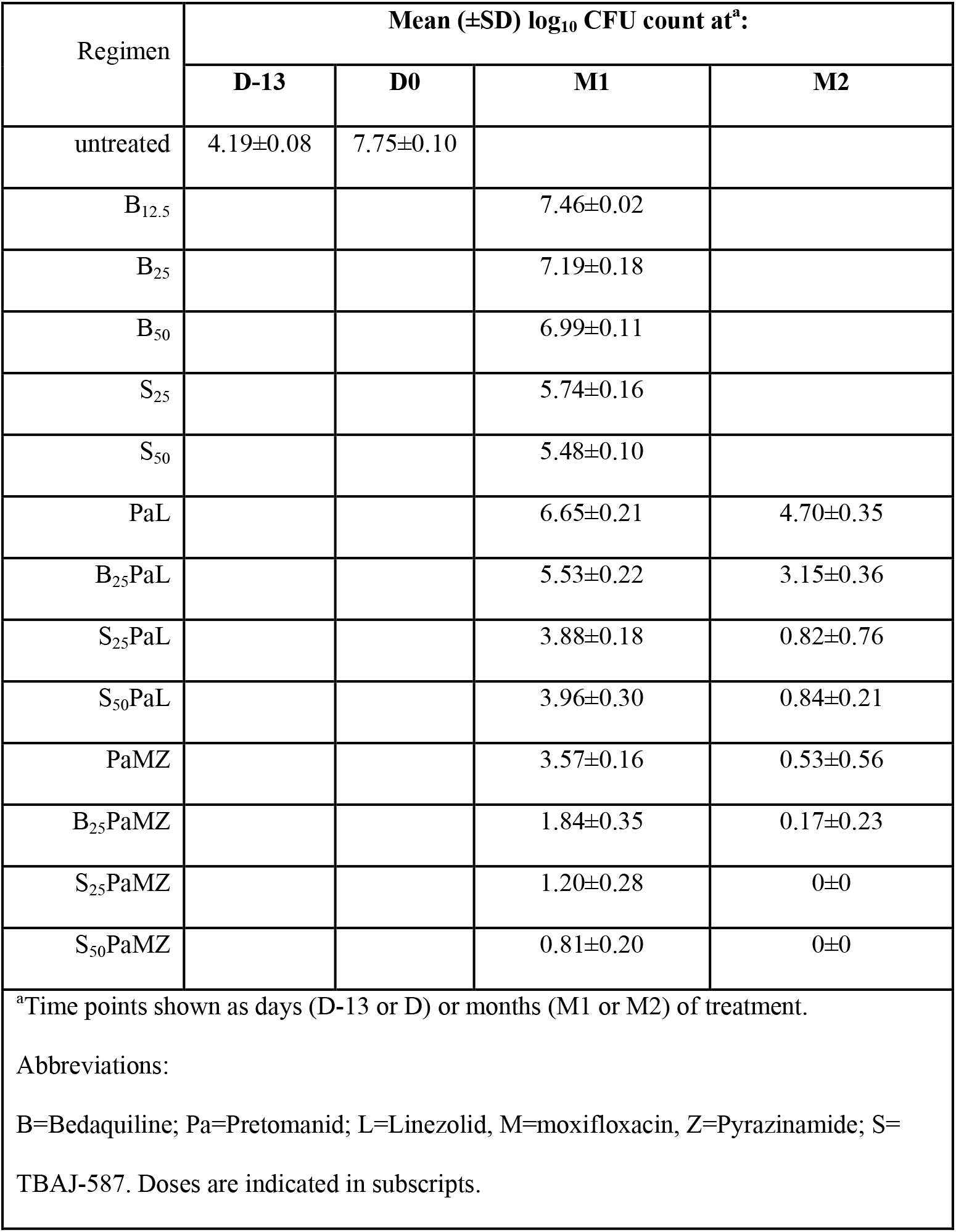
Lung CFU counts assessed during treatment against the *M. tuberculosis* H37Rv strain with an *Rv0678* mutation.

